# Disrupted PDZ-domain binding motif of the dopamine transporter uniquely alters nanoscale distribution, dopamine homeostasis and reward motivation

**DOI:** 10.1101/2021.01.06.425643

**Authors:** Gunnar Sørensen, Mattias Rickhag, Damiana Leo, Matthew D. Lycas, Pernille Herrstedt Ridderstrøm, Pia Weikop, Freja Herborg, David Woldbye, Gitta Wörtwein, Raul R. Gainetdinov, Anders Fink-Jensen, Ulrik Gether

## Abstract

The dopamine transporter (DAT) is part of a presynaptic multi-protein network involving interactions with scaffold proteins via its C-terminal PDZ-domain binding sequence. In a mouse model expressing DAT with mutated PDZ binding sequence (DAT-AAA), we previously demonstrated the importance of this binding sequence for striatal expression of DAT. Here we show by application of direct Stochastic Reconstruction Microscopy (dSTORM) not only that the striatal level of transporter is reduced in DAT-AAA mice, but also that the nanoscale distribution of the transporter is altered with a higher propensity of DAT-AAA to localize to irregular nanodomains in dopaminergic terminals. In parallel, we observe mesostriatal dopamine (DA) adaptations and changes in DA-related behaviors different from those seen in other genetic DAT mouse models. DA levels in striatum are reduced to ∼45% of wild type (WT), accompanied by elevated DA turnover. Nonetheless, Fast-Scan Cyclic Voltammetry recordings on striatal slices reveal a larger amplitude and prolonged clearance rate of evoked DA release in DAT-AAA mice compared to WT mice. Autoradiography and radioligand binding show reduced DA D2 receptor levels while immunohistochemistry and autoradiography show unchanged DA D1 receptor levels. In behavioral experiments, we observe enhanced self-administration of liquid food under both a fixed-ratio (FR1) and progressive-ratio (PR) schedule of reinforcement, but a reduction compared to WT when using cocaine as reinforcer. Summarized, our data demonstrate how disruption of PDZ-domain interactions causes changes in DAT expression and its nanoscopic distribution that in turn alter DA clearance dynamics.

---

Dopamine (DA) plays a fundamental role as a modulatory neurotransmitter controlling locomotion, reward, cognition and endocrine functions. Aberrant dopaminergic signaling is associated with neuropsychiatric diseases such as schizophrenia, attention-deficit hyperactivity disorder (ADHD) and drug addiction (1-3). The presynaptic DA transporter (DAT) is responsible for clearing released DA from the extracellular space and represents the major target for psychostimulants, such as cocaine and amphetamine (4-6) Importantly, the generation and characterization of genetically modified mice and rats with altered DAT expression has supported a pivotal role of DAT in regulating DA homeostasis. Mice and rats lacking DAT (DAT knock-out; DAT-KO) and mice with reduced DAT expression (DAT knock-down; DAT-KD) (7-9) exhibit profound alterations in extracellular DA dynamics with impaired DA clearance that lead to adaptive processes in the pre- and postsynaptic compartments (10,11). The alterations in DA dynamics is, moreover, accompanied by critical behavioral changes (5,7-9). Profound changes in DA dynamics have also been observed in knock-in mice expressing neuropsychiatric disease-associated DAT mutations (12,13)

It is well established that the DA system, and in particular DA neurons projecting from the ventral tegmental area (VTA) to the nucleus accumbens, plays an essential role in addiction to drugs, such as cocaine, methamphetamine, heroin, and nicotine, that elicit major increases in extracellular nucleus accumbens DA levels through distinct pharmacological mechanisms (14). Abuse of alcohol and food has been linked to DA release as well, further supporting that shared mechanisms are likely to underlie the compulsive reward-seeking behavior seen in addicted individuals (14). In addition, there is compelling evidence suggesting that abuse promotes common adaptive changes in the DA system. A key finding in human studies, for example, is a marked decrease in striatal D2 receptor (D_2_R) binding, which is believed to result from repetitive and long-lasting stimulation of this high-affinity receptor leading to internalization and receptor down-regulation (15-19). Indeed, the reduced D_2_R levels have been associated with a propensity for impulsive and compulsive behaviors and thus to be an important biomarker for the addicted brain (14). A decrease in the vesicular storage of pre-synaptic DA has furthermore been observed in human addicts, representing yet another key biological feature (19-23). However, although it is evident that alterations in D_2_R binding and striatal DA release might constitute interesting surrogate markers for the addicted brain (24-26), we still have a poor understanding of the mechanisms underlying these adaptations of the DA system and their precise biological effects.

We have generated a DAT knock-in mouse strain (DAT-AAA) with modified C-terminus to investigate the significance of PDZ (PSD-95/Dlg/ZO-1) domain scaffold interactions for DAT function and DA homeostasis (27). In these mice, the C-terminal PDZ-target sequence (−LLV) has been substituted for three alanines (−AAA), resulting in abolished PDZ-domain mediated interactions that cause extensive loss of striatal DAT levels down to 10-20 % of that seen in wild-type (WT) mice (27). Our data suggested that this loss is not caused by impaired folding and maturation of the protein but rather by enhanced degradation likely involving increased constitutive internalization of the transporter (27). An interesting question is whether this cellular phenotype also involves alteration in the nanoscopic distribution of the transporter. Indeed, our previous studies using super-resolution dSTORM have revealed that in both cultured and brain slices DAT is found to be sequestered into discrete PIP2 (phosphatidylinositol-4,5-bisphosphate)-enriched nanodomains in the plasma membrane of presynaptic varicosities and neuronal projections (28,29). These nanodomains were proposed to enable the neuron to rapidly move the transporter between different functional localizations and thereby optimize availability and activity of the transporter at nanoscale levels in the dopaminergic terminals (28,29). Here, we apply super-resolution dSTORM to assess the nanoscale distribution of DAT in DAT-AAA mice as compared to WT mice in striatal slices. Importantly, the data show that both the WT and the DAT-AAA mutant are found in discrete nanodomains in dopaminergic terminals of the slices. The data moreover indicate that, despite lower expression, The DAT-AAA mutant is more prone to exist in these nanodomains than the WT transporter with a higher fraction of nanodomains >75nm in diameter. Of major interest, we find that this altered nanoscale distribution of DAT-AAA, together with the decreased expression, is accompanied by unique homeostatic changes to the dopaminergic system that are distinct from those seen in other DAT genetic mouse models with reduced DA uptake capacity. We observe accordingly that, although the striatal DA levels are reduced in DAT-AAA mice to 45% of WT with a parallel increase in DA turnover, the amplitude of evoked DA release, as assessed by fast-scan cyclic voltammetric (FSCV) recordings from striatal slices, is markedly higher in DAT-AAA mice alongside with a prolonged clearance rate. Furthermore, we observe significant reduction in striatal D_2_R binding and striking changes in motivational behavior including enhanced self-administration of liquid food but decreased self-administration of cocaine as compared to WT mice. In summary, the data substantiate the critical importance of presynaptic PDZ-domain mediated protein-protein interactions for proper temporal and spatial regulation of striatal DA homeostasis by DAT.

## Results

### Altered nanoscale distribution of DAT in DAT-AAA mice

We have recently shown by application of dSTORM that DAT distributes into cholesterol-dependent and PIP2-enriched nanodomains in the plasma membrane of presynaptic varicosities and neuronal projections of dopaminergic neurons (28,29). To assess whether this nanoscale distribution of DAT was regulated by PDZ domain scaffold protein interactions, we decided to visualize DAT by dSTORM in striatal slices obtained from DAT-AAA mice and WT littermates. To identify dopaminergic terminals, we stained as well for the vesicular monoamine amine transporter 2 (VMAT2). The resulting reconstructed dSTORM images showed numerous irregular dopaminergic terminals/varicosities in slices from WT mice with both DAT and VMAT2 signal (Fig. 1A, top, example highlighted by yellow arrows). In slices from DAT-AAA mice, the number of DAT localizations was decreased, as expected, while the density and distribution of VMAT2 localizations were similar to that seen in the WT (Fig. 1, middle, example highlighted by yellow arrows). To further assess the distribution in the dopaminergic terminals, we filtered the dSTORM images for likely dopaminergic presynaptic sites through an adaptation of Voronoi tessellation (30,31) that we recently developed when analyzing DAT distribution in WT slices (29). In brief, DAT and VMAT2 signals were combined, and structures that were in the size range of varicosities and made up from localizations of a required density were isolated. By utilizing combined data from DAT and VMAT2 signals into this algorithm, presynaptic varicosities in both WT and DAT-AAA slices could be located (Fig. 1A, bottom). Dopaminergic presynaptic sites in the WT slices were often distinct by the occurrence of DAT in a clustered distribution adjacent to the VMAT2 signal (Fig 1A), similar to what we have observed before in cultured DA neurons (28). In the DAT-AAA slices, there were far fewer DAT localizations compared to WT present on the presynaptic varicosities, however, the signal appeared to be highly clustered and often adjacent to the VMAT2 signal (Fig. 1A, bottom). Analysis of the data by Density-Based Spatial Clustering of Applications with Noise (DBSCAN) (28,32) substantiated clustering of DAT into irregular nanodomains in dopaminergic terminals from both WT and DAT-AAA mice (Fig. 1B, C). We decided to analyze the clusters in two ways. First, we assessed a possible difference in cluster size by arbitrarily calculating clusters that were larger than 75 nm in diameter. Interestingly, despite the reduced level of DAT in DAT-AAA slices, there was a greater proportion of large (>75 nm in diameter) transporter clusters for DAT-AAA as compared to WT (Fig 1B). This was particularly evident on apparent varicosities with fewer localizations (Fig 1B). Second, we determined the fraction of DAT localizations present in dense clusters, which not surprisingly showed that overall, there was a greater fraction of WT DAT localizations found in dense clusters (Fig. 1C). This likely reflected the increased amount of DAT found on a given varicosity because when separated by the number of DAT localizations found on a given varicosity, there were more DAT-AAA localizations found within dense clusters than in comparable WT varicosities (Fig 1C). That is, the fraction of localizations in nanodomains was higher at comparable densities between DAT-AAA and WT (red dots versus black dots, Fig. 1B), indicating a higher propensity for DAT-AAA to be present within a nanodomain. Thus, disruption of the C-terminal PDZ binding sequence of DAT does not only dramatically reduce expression of the transporter but also leads to increased clustering into nanodomains of the remaining DAT in dopaminergic terminals of the striatum.

**Figure 1.**
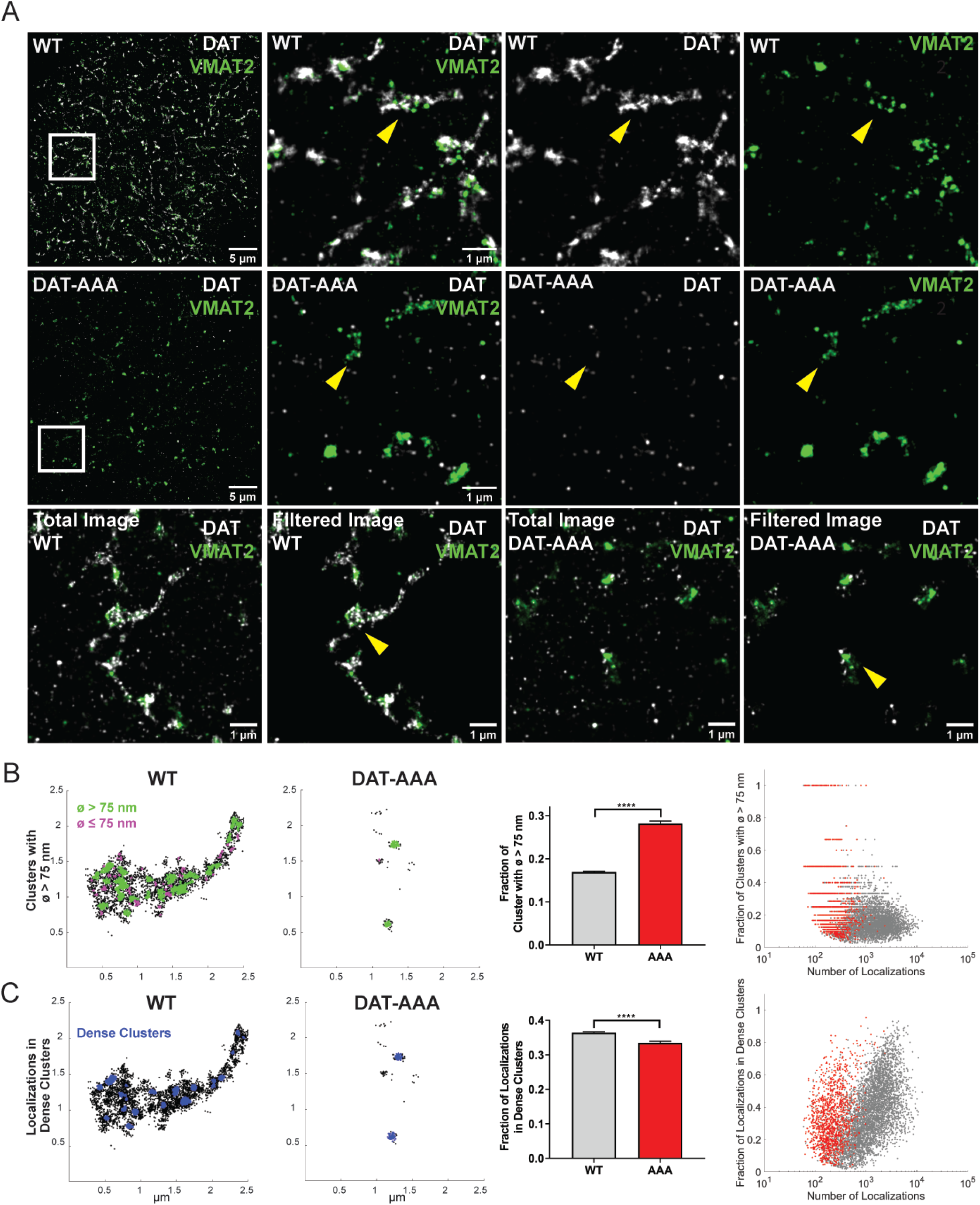
Visualization of DAT by super-resolution dSTORM imaging in striatal slices shows evidence for a hyperclustered phenotype of DAT-AAA. *A*, Example dual color dSTORM images for DAT (white) and for VMAT2 (green) from the striatum of a WT mouse and a DAT-AAA mouse. *Top panel*, WT mouse striatial section (left) with region of interest (white box) shown in the other images: middle left, merged DAT and VMAT2 signal; middle right, DAT signal; right, VMAT2 signal. *Middle panel*, DAT-AAA mouse striatial section (left) with region of interest (white box) shown in the other images: middle left, merged DAT and VMAT2 signal; middle right, DAT signal; right, VMAT2 signal.*Lower panel*, Example WT and DAT-AAA images before (left and middle right) and after filtering (middle left and right) for dopaminergic terminals/varicosities (see methods). Yellow arrows indicate examples of presumed dopaminergic terminals/varicosities. *B*, Example WT varicosity (left) and DAT-AAA varicosity (middle left), where clusters were identified with DBSCAN parameters 15 nm search radius and 5 number of points. Clusters with diameter >75 nm are colored in green and remaining clusters are colored in magenta. Despite fewer DAT localizations for DAT-AAA, there is a larger fraction of clusters with diameter > 75 nm. Quantification of the data are shown as fraction of nanodomains >75 nm (mean±S.E., unpaired t-test P <0.0001) (middle panel) and as a comparison of the fraction of clusters identified with a diameter > 75 nm and the number of localizations found in each detected varicosity (WT, grey dots; DAT-AAA red dots). *C*, Example WT varicosity (left) and DAT-AAA varicosity (middle left) comparing number of DAT localizations per varicosity that are present in dense clusters (identified with DBSCAN parameters 25 nm search radius and 30 number of points, shown in blue). Quantification of the data are shown as fraction of localizations found within large dense clusters (mean±S.E., unpaired t-test P <0.0001) (middle panel) and as a comparison of the number of localizations found in each detected varicosity with the fraction of DAT localizations found within large dense clusters. Despite fewer localizations for DAT detected in the DAT-AAA varicosities (red dots), more large dense clusters were observed than in WT varicosities (grey dots) with equivalent number of localizations. Data are from 4473 WT varicosities and 1186 DAT-AAA varicosities from 30 dSTORM images taken from 3 animals for each condition.

### Decreased DA tissue content in DAT-AAA mice

Next, we assessed possible alterations in DA homeostasis occurring as a result of the reduced striatal DAT expression together with the apparent change in nanoscale distribution of DAT in DAT-AAA mice as described above. We determined total striatal tissue content of DA and DA metabolites (Fig. 2A). The analysis of the striatal homogenates revealed a decrease in total DA levels in the DAT-AAA mice to ∼45% of WT (WT, 3.8±0.3 µg/g tissue; DAT-AAA, 1.68±0.14 µg/g tissue, p<0.0001), but increased homovanillic acid (HVA) concentrations (WT, 0.71±0.04 µg/g tissue; DAT-AAA, 1.48±0.12 µg/g tissue, p<0.0001) as compared to WT. There was no difference in 3,4-dihydroxyphenylacetic acid (DOPAC) levels (WT, 0.73±0.12 µg/g tissue; DAT-AAA, 0.6±0.07 µg/g tissue, p>0.05, Fig. 2A). However, both the HVA/DA ratio (∼4.8-fold, P<0.0001) and the DOPAC/DA ratio (∼1.8-fold, p<0.05) were significantly increased, indicating increased DA-turnover in DAT-AAA mice (Fig. 2B).

**Fig. 2.**
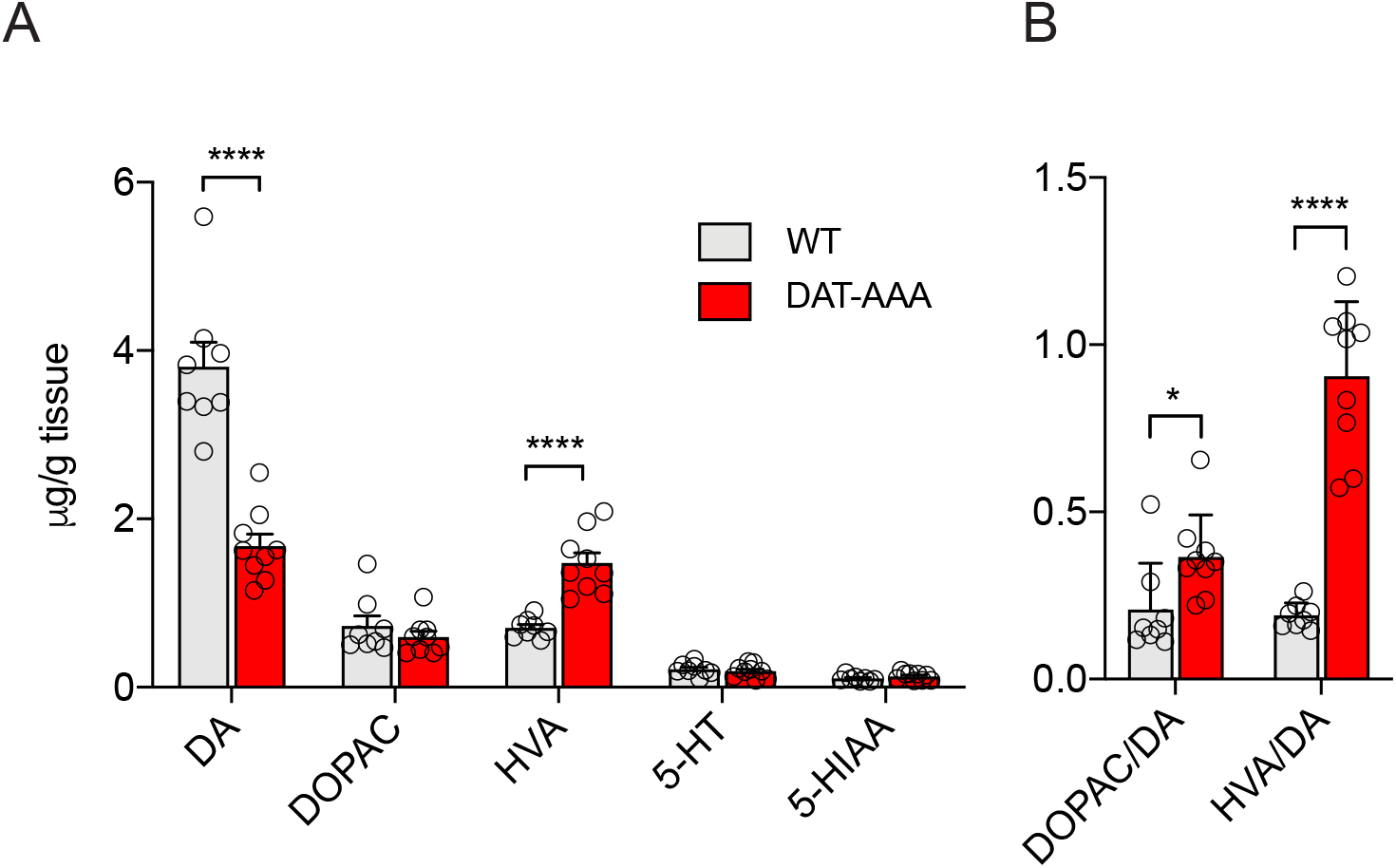
Striatal tissue content of DA and metabolites. *A*, Total tissue content of DA and major metabolites in striatal homogenates from DAT-AAA and WT mice. The content of DA was significantly decreased (∼45% of WT) in the DAT-AAA mice while the content of HVA was significantly increased (∼2-fold). Data are means ± S.E, WT: n=8, DAT-AAA: n=9), ****p<0.0001, DAT-AAA vs. corresponding WT group, Holm-Sidak post-hoc test after two-way ANOVA (significant effect of metabolite and genotype, F_(4,75)_=165.7, p<0.0001 and F_(1,75)_=15.93, p=0.0002, respectively). B, DOPAC/DA and HVA/DA turnover ratios. DAT-AAA animals displayed a 1.8-fold increased DOPAC/DA ratio indicative of increased DA-synthesis, but also 4.8-fold increased HVA/DA ratios indicative of an increased turnover of DA. Data are means ± S.E, WT: n=8, DAT-AAA: n=9, *p<0.05, ****P<0.001, Holm Sidak post-hoc test after two-way ANOVA (significant effect of ratio and genotype, F_(1,30)_=25.97, p<0.0001 and F_(1,30)_=72.28, p<0.0001, respectively).

### Enhanced evoked release and prolonged clearance of DA in DAT-AAA mice

To investigate the dynamics of vesicular DA release and reuptake, we performed FSCV recordings on striatal brain slices. Interestingly, in the slices derived from DAT-AAA mice we observed elevated peak concentrations of DA (∼7.5 fold higher than WT) after stimulated release (WT, 0.072±0.025 µM; DAT-AAA, 0.544 ±0.27 µM, p<0.05) (Fig. 3A, B). Consistent with the markedly reduced DAT levels, the data moreover revealed substantially prolonged clearance (∼4 fold longer) of DA in DAT-AAA mice as compared to WT (WT, tau=0.62±0.06; DAT-AAA, tau= 2.4±2.8, p<0.05, Fig. 3C).

**Figure 3.**
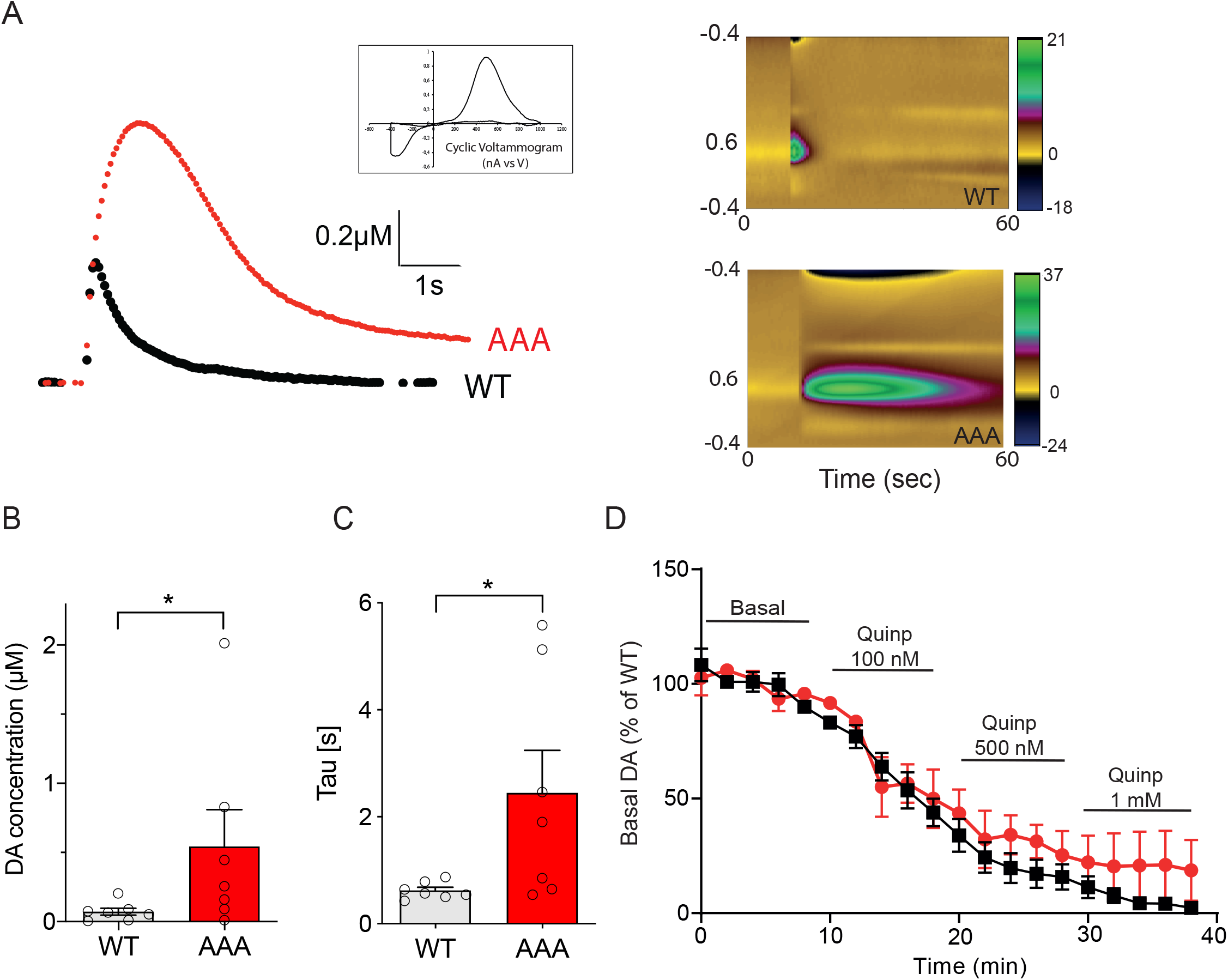
Enhanced vesicular DA release and prolonged clearance rate of extracellular DA in DAT-AAA mice. *A*, DAT-AAA mice show an increase in evoked DA release in striatal slices as measured by Fast-Scan Cyclic Voltammetry. *Left*, Representative stimulated DA-release shown (DAT-AAA, red; WT, black), *Right*, Color plots displaying sequential voltammograms indicating larger and more prolonged increase in DA levels for DAT-AAA (lower panel) than for WT (upper panel) (time in sec, x axis; applied voltage, y axis; z axis; measured current in color). *B*, Peak concentrations of DA following stimulated response in striatal slices show elevated DA levels. Data are means ± S.E. of measurements on 45 slices from 7 WT mice and on 55 slices from 7 DAT-AAA mice; * p<0.05, Wilcoxon matched-pairs signed rank test. C, Clearance of DA is prolonged in DAT-AAA mice. Data are presented as tau, * p<0.05, Student’s t-test. *D*, Dose-response treatment using the D_2_R-agonist quinpirole show blockade of stimulated DA release as measured by fast-scan cyclic voltammetry in striatal slices. Data are means ± S.E, WT, n=10; DAT-AAA, n=7.

We next addressed possible changes in presynaptic D_2_R autoreceptor function as impaired auto-receptor function could contribute to the elevated levels of released DA in the DAT-AAA mice. Accordingly, we determined by FSCV the effect of the D_2_R agonist quinpirole on stimulated DA release. In both DAT-AAA mice and WT mice, quinpirole blocked DA release, supporting preserved D_2_R autoreceptor function in DAT-AAA mice (Fig. 3D).

### Unaltered D_1_R immunoreactivity and binding sites in DAT-AAA mice

To investigate the cellular/regional distribution of the DA D1 receptor (D_1_R) in DAT-AAA and WT mice, we determined D_1_R-immunoreactivity (D_1_R-ir) in the mesostriatal dopaminergic system. Dense D_1_R-ir was observed in dorsal/ventral striatum, substantia nigra and ventral tegmental area as well as olfactory tubercles. DAT-AAA mice showed similar intensities of D_1_R-ir in both striatal and midbrain areas compared to WT mice, thus suggesting unaltered expression of the receptor (Fig. 4A). In midbrain, D_1_R distribution was predominant in the substantia nigra pars reticulata but again no difference was observed between genotypes (Fig. 4B). Importantly, the observed D_1_R-distribution was in accordance with earlier reports (33).

**Figure 4.**
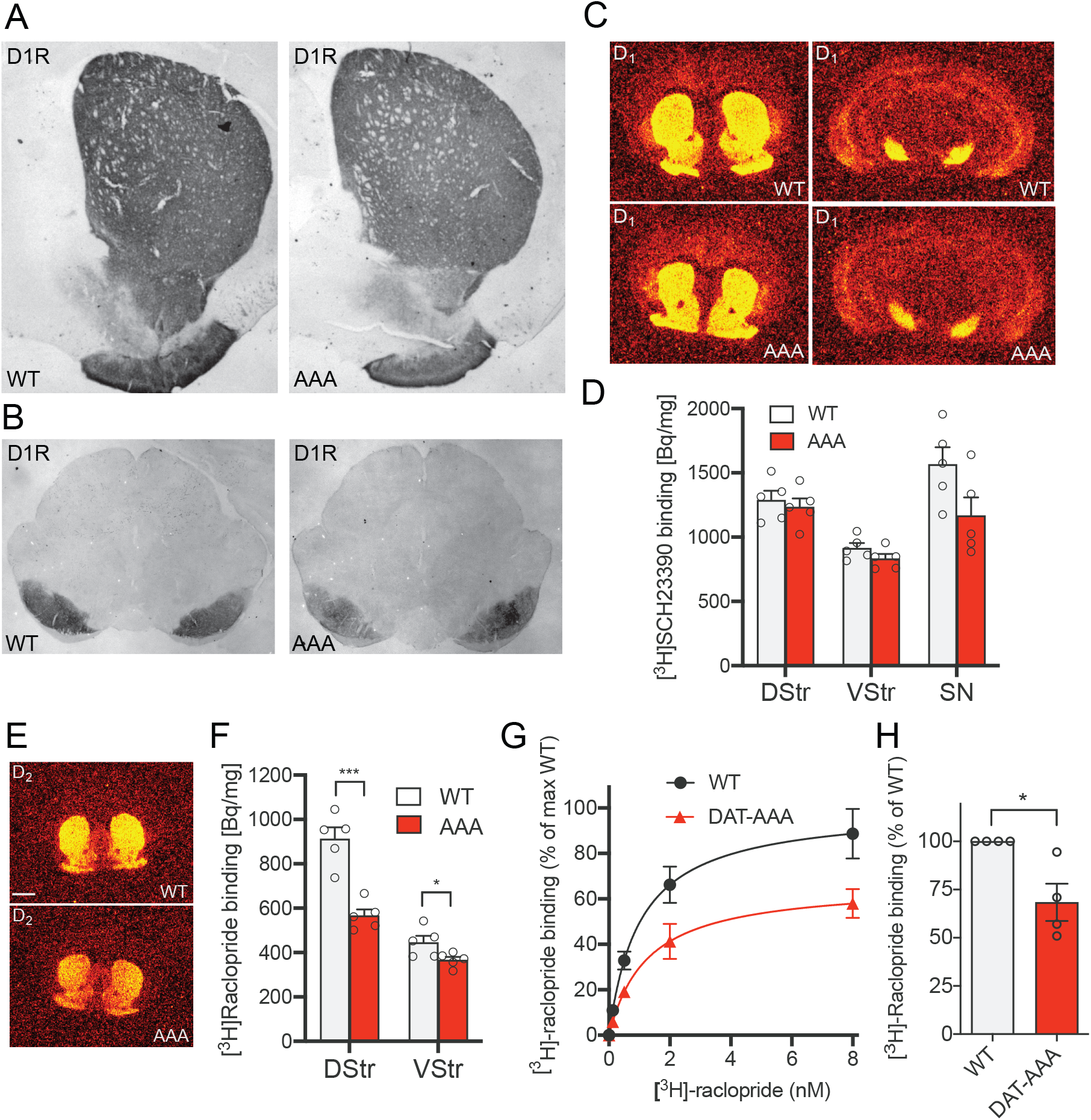
DA receptor expression and binding sites in DAT-AAA and WT mice. *A, B*, Representative photomicrographs show similar D_1_R immunoreactivity (D_1_R-ir) in striatum of WT and DAT-AAA mice (*A*) and ventral midbrain (*B*). In striatum, dense D_1_R-ir was observed particularly in the dorsal part while less D_1_R-ir was seen in ventral striatum. *C, D*, Assessment of functional D_1_R-binding sites were investigated using quantitative autoradiography. [^3^H]-SCH23390, a selective D_1_R-ligand, revealed a high density of D_1_R binding sites in the striatum and midbrain with similar intensities in both genotypes. Data are specific [^3^H]-SCH23390 binding in Bq/mg, means ± S.E, *n*=5 (dorsal striatum: WT, 1288±73 Bq/mg tissue; DAT-AAA, 1234±67 Bq/mg tissue, p=0.60; ventral striatum: WT, 914±40 Bq/mg tissue; DAT-AAA, 833±37 Bq/mg tissue, p=0.32, midbrain: WT, 1565±134 Bq/mg tissue; DAT-AAA, 1168±141 Bq/mg tissue, p=0.21, multiple t tests with Holm-Sidak correction). *E, F*, Assessment of functional D_2_R-binding sites were investigated using quantitative autoradiography. [^3^H]-Raclopride, a selective D_2_R ligand, revealed high density of D_1_R binding sites in the striatum of both WT and DAT-AAA mice, however, there was a significant reduction in binding in both the dorsal (DStr) and ventral striatum (VStr) of DAT-AAA mice. Data are [^3^H]-Raclopride binding in Bq/mg, means ± S.E, *n*=5 (dorsal striatum: WT, 912±52 Bq/mg tissue; DAT-AAA, 566±29 Bq/mg tissue, ***p<0.001; ventral striatum: WT, 446±30 Bq/mg tissue; DAT-AAA, 366±14 Bq/mg tissue, *p<0.05, multiple t tests with Holm-Sidak correction). No binding could be detected in the midbrain. *G*, Saturation radioligand binding experiments on striatal membranes using [^3^H]-raclopride, a D_2_R antagonist. Compiled normalized saturation curves demonstrate significantly reduced number of binding sites in striatum from DAT-AAA mice compared to WT. Data are means ± S.E, *n*=4 in % of the B_max_ for WT (206 ±25 fmol/mg protein). Equilibrium dissociation constant (K_d_) was not significantly different between the genotypes (K_d_: WT, 1.12 ±0.19 nM; DAT-AAA, 1.34 ±0.14 nM (*n*=4). *H*, B_max_ for DAT-AAA in % of B_max_ for WT. Data are means ± S.E, *n*=4, *p<0.05, one-sample t test.

In addition to immunohistochemical localization of the D_1_R, we assessed functional binding sites in striatum using quantitative autoradiography. Binding of the tritiated D_1_R antagonist [^3^H]-SCH23390 was determined in sections from both WT and DAT-AAA mice revealing a high density of D_1_R binding sites in dorsal and ventral striatum as well as in the midbrain with no difference between the genotypes (dorsal striatum: WT, 1288±73 Bq/mg tissue; DAT-AAA, 1234±67 Bq/mg tissue, p=0.60; ventral striatum: WT, 914±40 Bq/mg tissue; DAT-AAA, 833±37 Bq/mg tissue, p=0.32, midbrain: WT, 1565±134 Bq/mg tissue; DAT-AAA, 1168±141 Bq/mg tissue, p>0.21, Fig. 4C, D).

### Reduced DA D_2_-receptor binding sites in the striatum of DAT-AAA mice

We also performed quantitative autoradiography for D_2_R by using the tritiated D_2_R antagonist [^3^H]-raclopride. In WT mice, we observed a high density of D_2_R binding sites in dorsal and ventral striatum. In DAT-AAA mice, however, [^3^H]-raclopride binding was reduced in both regions with the most prominent reduction in the dorsal striatum (dorsal striatum: WT, 912±52 Bq/mg tissue; DAT-AAA, 566±29 Bq/mg tissue, p<0.0001; ventral striatum: WT, 446±30 Bq/mg tissue; DAT-AAA, 366±14 Bq/mg tissue, p<0.04, Fig. 4E and F). Of note, we were unable to detect any specific [^3^H]-raclopride binding in the midbrain.

To substantiate the reduced binding of [^3^H]-raclopride in the striatum, we examined the density of D_2_R in striatal membranes in a [^3^H]-raclopride saturation binding experiment (Fig. 4G). In full agreement with the autoradiography data, the maximal number of binding sites (B_max_) was significantly reduced in DAT-AAA compared to WT mice (B_max_, 68±10% of WT, p<0.05) (Fig. 4H) without altered K_d_ (K_d_ DAT-AAA, 1.34±0.14 nM; K_d_ WT, 1.12±0.19 nM, p>0.05).

### Increased liquid food self-administration in DAT-AAA mice

Next, we wanted to investigate how the altered DA homeostasis in DAT-AAA mice affected DA-related behaviors. Because reduced D_2_R levels (15-19) and a reduced pool of DA (19-23) have been strongly associated with impulsive and compulsive behaviors related to addiction including binge eating, naïve DAT-AAA and WT mice were evaluated in a palatable liquid food self-administration paradigm. The self-administration was assessed both under a fixed ration (FR1) and progressive ratio (PR) schedule of reinforcement (Fig. 5). Under the FR1 schedule, both genotypes demonstrated liquid food intake that increased significantly with increasing concentrations of palatable liquid food (Fig. 5A). There was a significant effect of both food concentration and genotype (two-way ANOVA F_(4,65)_=20.35, p<0.0001 and F_(1,65)_=12.90, p=0.0006; p<0.001, respectively). Notably, post-hoc analysis showed an increased response rate in DAT-AAA mice at 10 % of liquid food compared to WT mice (p<0.05). Under the PR schedule, there was also a significant effect of palatable liquid food concentration (two-way ANOVA (F_(4, 60)_=14.53; p<0.001) and significant effect of genotype (F_(1, 60)_=38.94; p<0.001) with no genotype by food concentration interaction (F_(4, 60)_=0.54, p=0.70). Post-hoc analysis showed that DAT-AAA mice reached significantly higher breaking points at 3, 10, 32 and 100% of liquid food compared to WT mice (p<0.05 and p<0.01) (Fig. 5B). Together, the data show that DAT-AAA mice are willing to work harder for a food reward than WT mice.

**Figure 5.**
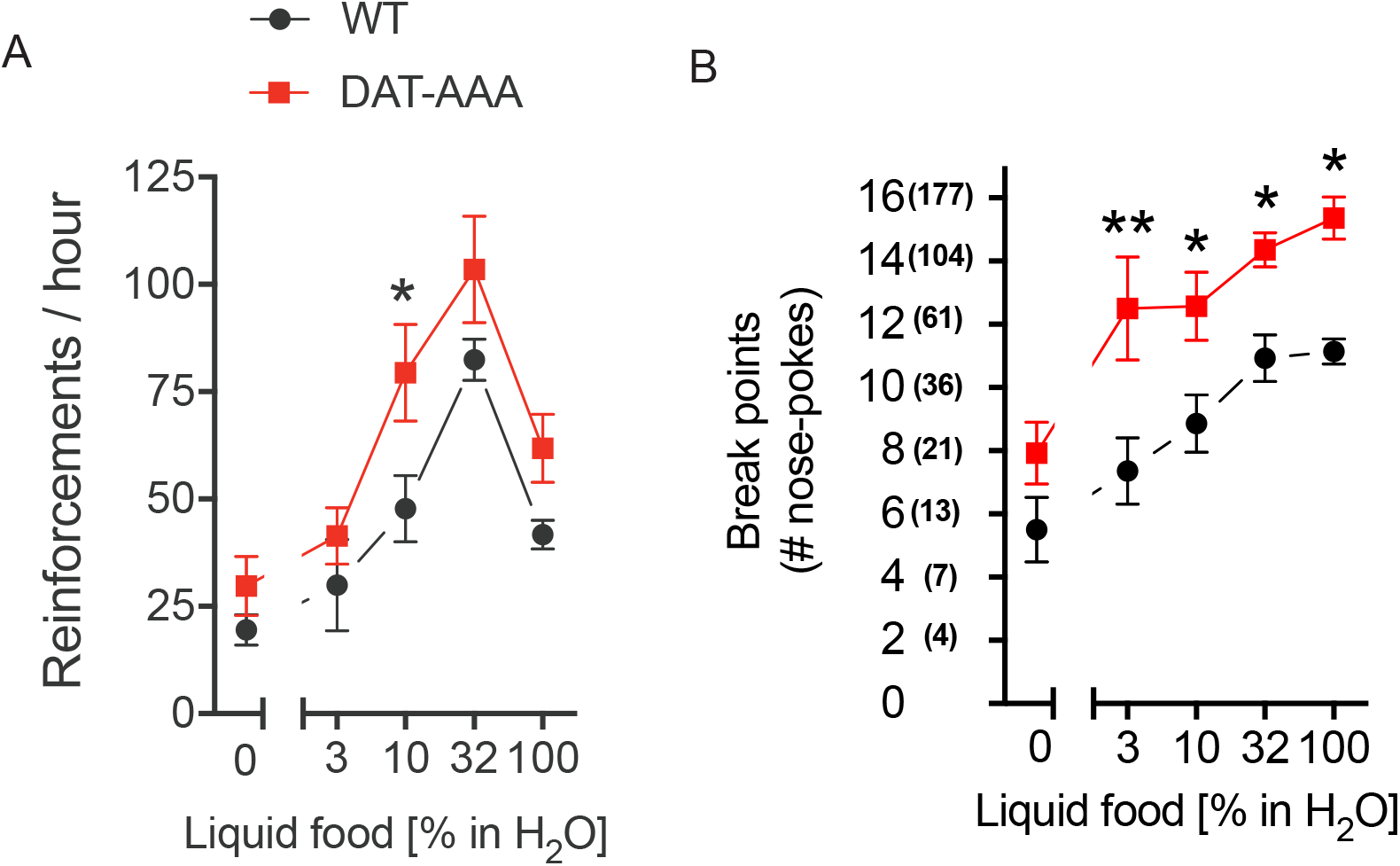
Operant self-administration of liquid food reinforcements. Food-maintained operant behavior under the FR 1 (A) and the PR (B) schedule of reinforcement in DAT-AAA and WT. A, DAT-AAA showed a higher number of nose pokes than WT mice in food-maintained responding under the FR 1 schedule. Data are means ± S.E., WT n=7, DAT-AAA, n=8, *p<0.05, DAT-AAA vs WT group, Holm-Sidak post-hoc test following two-way ANOVA (significant effect of food concentration and genotype, F_(4,65)_=20.35, p<0.0001 and F_(1,65)_=12.90, p=0.0006, respectively). B, Under the PR schedule of reinforcement, DAT-AAA mice reached higher breaking points at a concentration of 3, 10, 32 and 100% liquid food. Data are means ± S.E, n=7, ** p<0.01, * p<0.05, DAT-AAA vs WT group, WT; Holm-Sidak post-hoc test following two-way ANOVA (significant effect of liquid food concentration and genotype, F_(4,60)_=14.53; p<0.001 and F_(1,60)_=38.94; p<0.001, respectively, with no genotype by food concentration interaction, F_(4, 60)_=0.54, p=0.70).

### Attenuated cocaine self-administration in DAT-AAA mice

We next evaluated whether DAT-AAA mice also would show increased motivation to self-administer cocaine or whether the dramatic reduction in expression and distribution of its main target, DAT, would affect the pharmacological action of cocaine. The mice were tested under both FR1 and PR schedules. Strikingly, DAT-AAA mice did not self-administer cocaine (p=0.76) whereas WT mice did. Under the fixed ratio schedule, there was a significant effect of cocaine dose (two-way ANOVA F_(4,80)_=2.76; p=0.03) and a significant effect of genotype (F(_1,80)_=4.38; p=0.04) with no genotype by dose interaction (F(_4, 80)_=1.79, p=0.14). Post-hoc analysis indicated that DAT-AAA mice responded with significantly fewer nose-pokes at 0.32 mg/kg/infusion (p< 0.05) of cocaine compared to WT mice (Fig. 6A). Similarly, under the PR schedule of reinforcement, two-way ANOVA revealed a significant effect of cocaine dose (two-way ANOVA, F_(4,25)_=9.32; p<0.001) and a significant effect of genotype (F(_1,25)_=10.78; p=0.0030) with no genotype by dose interaction (F_(4, 25)_= 2.47 p=0.07) (Fig. 6B). Holm-Sidak post-hoc analysis showed that DAT-AAA mice responded with significantly fewer nose-pokes at 0.3.2 mg/kg/infusion (p<0.01) of cocaine compared to WT mice (Fig. 6B). Thus, the pharmacological effect of cocaine appeared clearly hampered in DAT-AAA mice.

**Figure 6.**
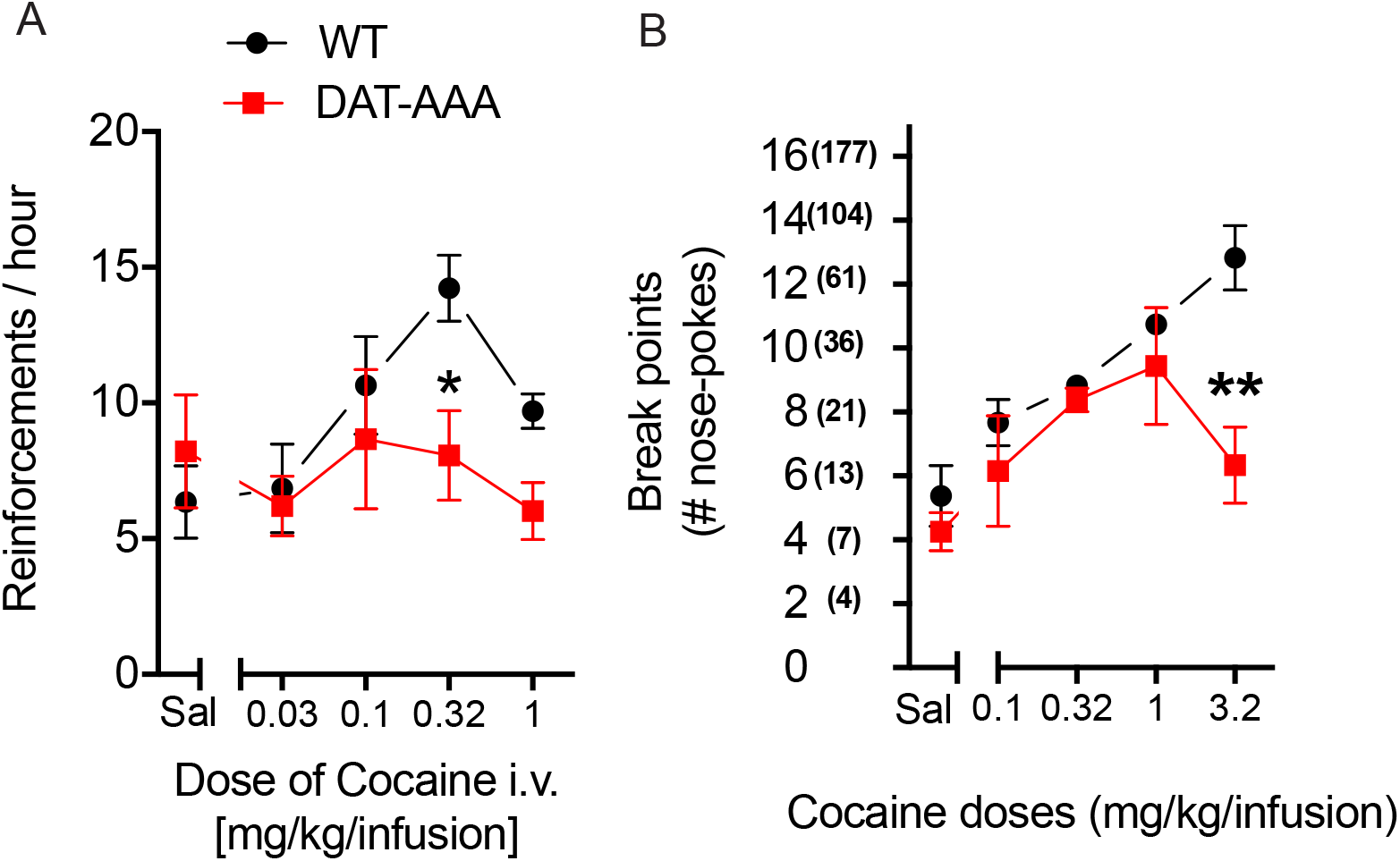
Operant self-administration of cocaine. Cocaine-maintained operant behavior under the FR 1 (*A*) and the PR (*B*) schedule of reinforcement in DAT-AAA and WT mice. *A*, DAT-AAA mice showed an attenuated response compared to WT mice in cocaine-maintained responding under the FR 1 schedule. Data are means ± S.E, n=9, * p<0.05, DAT-AAA vs WT, Holm-Sidak post-hoc test following two-way ANOVA (significant effect of cocaine dose and genotype, F_(4,80)_=2.76; *p*=0.03 and F_(1,80)_=4.38; *p*=0.04, respectively, with no genotype by dose interaction, F(_4, 80)_=1.79, p=0.14). *B*, DAT-AAA mice showed an attenuated response compared to WT mice in cocaine-maintained responding under the PR schedule. Data are means ± S.E, n=4, ** p<0.01, DAT-AAA vs WT, Holm-Sidak post-hoc test following two-way ANOVA (significant effect of cocaine dose and genotype, F_(4,25)_=9.32; p<0.001 and F_(1,25)_=10.78; p=0.0030, respectively, with no genotype by dose interaction, F_(4, 25)_= 2.47 p=0.07.

This was further supported by data from in vivo microdialysis experiments showing that administration of cocaine (30 mg/kg) resulted in an expected large increase in extracellular DA levels in WT mice. The increase peaked at 415±48% of basal level 40 min following injection before it gradually declined. In DAT-AAA mice, however, we observed an attenuated response to cocaine compared to WT mice with DA levels peaking at 184±24% of basal level 60 min after injection (Fig. 7A). A three-way repeated measures (RM) ANOVA showed a significant effect of genotype (F_(1, 12)_ = 48.76, p<0,0001), time (F _(9, 108)_ = 26.93, p<0,0001) and cocaine treatment (F_(1, 12)_ = 92,09, p<0,0001). Post hoc analysis showed that cocaine treatment was associated with a significant genotype effect (WT vs DAT-AAA) at multiple time points (20-100 min following cocaine treatment, p<0.01-0.0001, Fig. 7A). Overall, the data demonstrate that administration of a dose of cocaine (30mg/kg s.c.) elicited a 4-fold increase in extracellular DA levels in WT mice which was significantly attenuated in the DAT-AAA mice.

**Figure 7.**
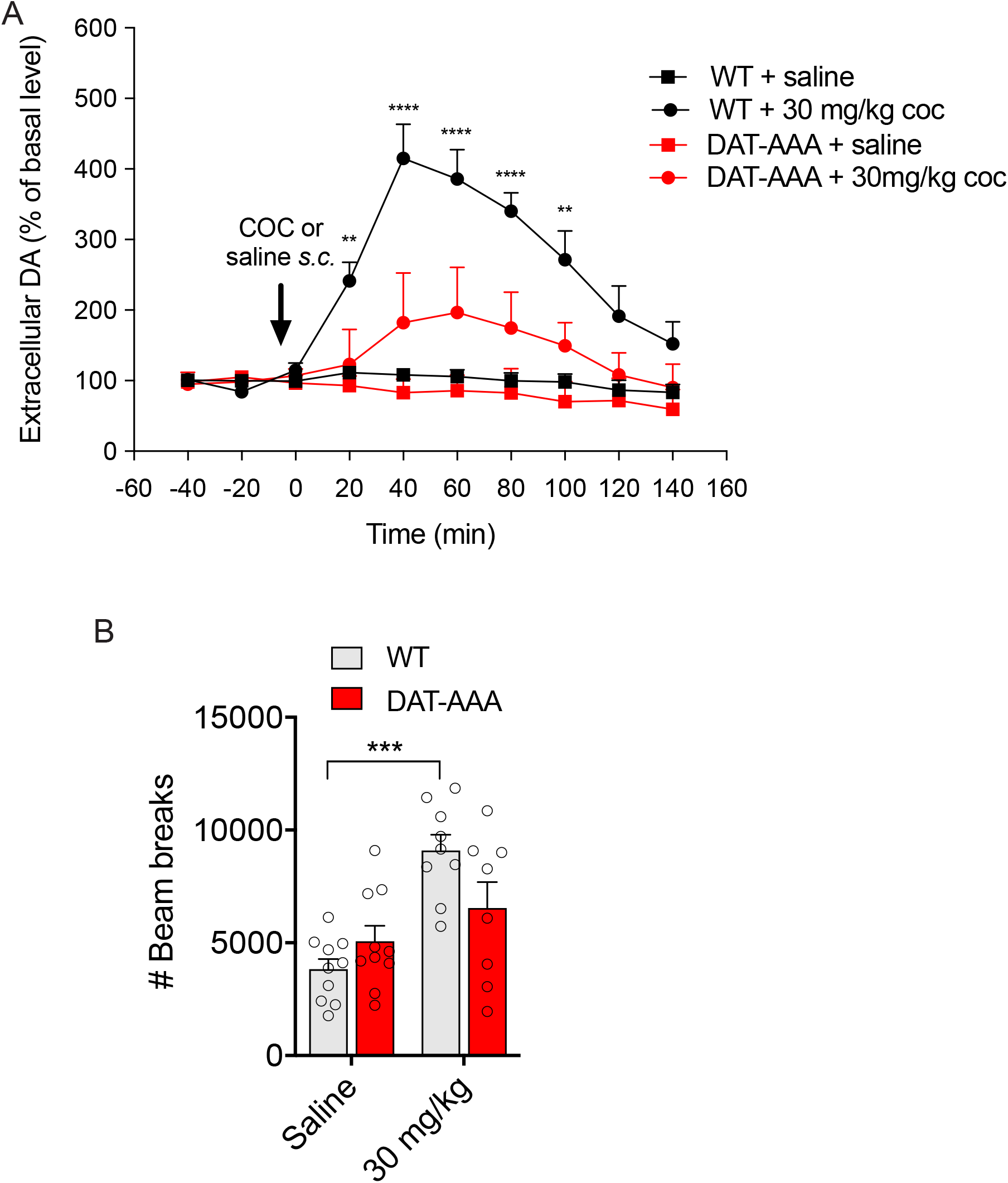
Effects of cocaine on extracellular dopamine levels and locomotion. *A, B*, Effect of cocaine (30 mg/kg *s*.*c*.) or saline on extracellular DA levels measured by microdialysis (20 min fractions) in the dorsal striatum of freely moving WT (*A*) or DAT-AAA mice (B). Extracellular DA levels increased 4-fold in WT mice after cocaine administration while DAT-AAA mice only displayed ∼1.8-fold increase in extracellular DA levels. Data are means±S.E., WT, n=6; DAT-AAA, n=8). Black and red bars represent ****p<0.0001 or ** p<0.01, of WT and DAT-AAA mice, respectively. ****p<0.001, ** p<0.01, genotype effect (WT vs DAT-AAA mice) after three-way ANOVA and Holm-Sidak post hoc test (cocaine treatment F _(1, 12)_ = 92.09, p<0,0001). C, Effect of cocaine on locomotor activity in DAT-AAA and WT mice. The 1-hour locomotion test shows impaired response to cocaine (30 mg/kg) in DAT-AAA as compared to WT. Data are means±S.E.; DAT-AAA, n=8; WT, n=9, ***p<0.001, cocaine (30 mg/kg i.p.) vs saline for WT (no significance for DAT-AAA, p=0.35), Holm-Sidak post hoc analysis hoc test after two-way ANOVA (significant effect of cocaine treatment, F(1, 33)=20.38; p<0.001, but no significant effect of genotype, F(1,33)=0.77; p=0.39.

We finally assessed whether the effect of cocaine on locomotor activity in DAT-AAA mice was affected as well. Indeed, the 1-hour locomotion test revealed that alanine substitutions of the C-terminal PDZ domain in DAT-AAA mice were associated with a dampened response to cocaine (significant genotype x dose interaction in a two-way ANOVA F_(1, 33)_=6.41; p<0.05; Figure 7B). A two-way ANOVA showed a significant effect of cocaine treatment (two-way ANOVA, F_(1, 33)_=20.38; p<0.001) but no significant effect of genotype (F(_1,33)_=0.77; p=0.39). Holm-Sidak post hoc analysis showed that cocaine (30 mg/kg i.p.) increased locomotor activity in WT mice (p<0.001) but did not have any effect in DAT-AAA mice compared to saline groups (Fig. 7B).

## Discussion

Presynaptic DAT sequesters released DA from the synapse and plays a major role in controlling DA homeostasis. We have previously reported that the C-terminal PDZ domain binding sequence of DAT is required for proper targeting and membrane stabilization of DAT in striatal terminals (27). Here, we show that distinct adaptations of the mesostriatal DA system may occur as a consequence of reduced striatal DAT levels concomitant with an altered nanoscopic distribution of the transporter. Thus, the data substantiate the unequivocal importance of preserved PDZ domain interactions for proper DAT function and distribution, while at the same time revealing striking changes to the DA system.

We have earlier reported by employment of super-resolution microscopy that DAT distributes into nanodomains in varicosities/boutons of DA neurons. Our previous data suggested that the nanodomains consist of 10s of DAT molecules and that DAT preferentially assumes an inward facing conformation within these domains (28,29) The data also suggested that localization of DAT to nanodomains was tightly regulated; that is, while DA D2 autoreceptor activity promoted nanodomain localization, both acute and sustained membrane depolarization caused dispersal of DAT nanodomains. It was therefore speculated that DAT nanodomain distribution represents a means by which DA neurons can temporally and spatially switch DAT between distinct functional states to enable optimal availability of the transporter (28,29). Such speculations are in line with recent findings for other neuronal membrane proteins, including ion channels and receptors for which dynamic sequestering into nanoclusters is believed to be central for cellular function (see e.g. (34)).

In slices from DAT-AAA mice, we found, as expected, that the dSTORM DAT signal in the individual terminals/varicosities was much lower as compared to WT, yet the signal was still highly clustered and distributed into nanodomains. According to our analysis of thousands of terminals/varicosities, allowing for comparison of varicosities with similar levels of DAT in both DAT-AAA mice and WT mice, it appeared that the DAT-AAA mutant showed a higher propensity to be localized in nanodomains and that the fraction of larger nanodomains (>75 nm) was higher in slices from DAT-AAA mice than from WT mice. This suggests that unclustered DAT, rather than clustered nanodomain-localized DAT, interacts with still unknown PDZ domain proteins, and that these interactions promote the localization of DAT outside of its nanodomain distribution. An interesting consideration would be that breaking these interactions would not only increase nanodomain localization but also the constitutive internalization and degradation that we have reported for DAT-AAA (28). Indeed, membrane microdomains have previously been suggested to serve as “hot spots” for DAT internalization (35). Thus, although the precise molecular processes remain to be determined, our data add strong support for the existence of still poorly understood nanoscopic regulatory mechanisms with biological implications for the spatial and temporal function of DAT and consequently for DA homeostasis.

The neuroadaptative changes in the DA system of DAT-AAA mice are, according to our data, different from those seen in other DAT mutant mouse models. We observed, using of FSCV recordings in striatal slices, that evoked DA release in slices from DAT-AAA mice resulted in higher DA peak values with much slower DA clearance as compared to slices from WT mice. This result is different from findings for DAT-KO mice, where evoked DA release also revealed slower clearance but DA peak levels that were substantially lower and corresponding to only ∼25% of the values observed for WT mice (7,8). We would argue, that the lower peak DA levels, as compared to WT mice, would be expected in DAT-KO mice having only ∼5% of the total striatal DA tissue level seen in WT mice (11). In contrast, DAT-AAA mice maintain a relatively large DA pool (∼45% of WT levels), which not only suggests that even rather low reuptake capacity is sufficient to maintain a reasonable DA pool but also explain the DA “overshoot” as compared to WT, i.e. a pool that is nearly half of that seen in WT combined with reuptake capacity only 10-20% of WT would be predicted to result in larger DA peak levels. In this context, it is somewhat surprising that FSCV experiments on slices from for DAT-KD with 10% residual uptake capacity show markedly reduced peak DA levels compared to WT (8). Thus, despite reported similar uptake capacities, DAT-KD and DAT-AAA mice display differences in DA homeostasis. It is conceivable that these differences are related to the nature of the two transgenic mouse models; that is, whereas the reduction in uptake in DAT-KD mice is a direct result of reduced transcriptional activity, the reduced capacity in DAT-AAA mice is the complex result of disrupted PDZ domain interactions, leading to altered nanoscopic distribution and turnover of the transporter. We can at the present stage mainly speculate on how exactly these changes affect spatiotemporal DAT function, yet they still provide a possible explanation for the apparent differences between the models.

In the DAT-AAA mice, we also observed interesting adaptive changes of the DA receptors and these appeared to differ from those found in DAT-KO and DAT-KD mice as well. In DAT-KO mice, mRNA analysis showed a decrease (∼50%) in expression of both D_1_R and D_2_R (7) whereas in DAT-KD mice no difference was found in post-synaptic D_1_R and D_2_R availability although evidence was obtained for ∼50% reduction in D_2_R autoreceptors (7,8). In DAT-AAA mice, however, we observed a selective decrease in D_2_R binding and no change in D_1_R levels. Note that D_2_Rs have higher DA affinity and are therefore believed to be more prone to down-regulation than D_1_Rs (36). The decrease in D_2_R binding is interesting given that the most commonly observed trait for the addicted brain is a selective decrease in striatal D_2_R availability as a presumed consequence of increased extracellular DA levels in human addicts of cocaine, methamphetamine, alcohol and heroin (18,19) as well as obese overeaters (17). These changes in striatal D_2_R or D_2_R mRNA are observed as well in animals with an increased compulsive behavior towards drugs (37-39) and “fast food” (40). In DAT-AAA mice, we moreover found, as discussed above, a decreased pool of DA, which is considered another putative biomarker of addiction found in cocaine and heroin addicted patients (21,41). Thus, the adaptive changes observed in DAT-AAA mice seem to partially mimic those seen in the brain of addicted individuals.

Our behavioral analysis of the DAT-AAA mice revealed several interesting features. First, we observed that DAT-AAA mice are willing to work harder for a food reward than WT mice. It has been suggested that over-eating promoted by high-calorie foods, or “fast food” is controlled through the same pathways as those responsible for the rewarding effect of drugs of abuse (42-44) and, thus, that binge eating of palatable food will, similar to drugs of abuse, cause an acute increase in striatal extracellular DA (45). At the same time, lowered D_2_R levels have been associated with accelerated appearance of compulsive-like consumption of palatable food (42-44). Thus, the lowered D_2_R levels combined with higher evoked DA levels in DAT-AAA mice might directly underlie the willingness of the mice to work harder for a food reward than WT mice. Our data are in line with findings on DAT-KD mouse, who showed higher food intake in their home cages and an increased wanting of sucrose (46). In contrast, DAT-KO strains showed an unaltered number of days to learn operant behavior for food (47) or no difference in FR 5 and PR schedules for 20 mg food-pellets (48) most likely reflecting DA homeostatic differences between mice having no transporter and mice with ∼20% of WT uptake capacity as the DAT-AAA mice.

Although DAT-AAA mice worked harder for a food reward as compared to WT mice, they did not show increased willingness to work for a cocaine reward. The most likely explanation is that cocaine’s effect is blunted by the dramatically reduced level of DAT leading to a “cocaine-like state” with impaired clearance and elevated DA peak concentrations plus decreased D_2_R levels. In that sense, the data substantiate DAT as a primary target for the stimulatory action of cocaine in agreement with previous experiments in DAT-KO mice and in mice expressing a DAT variant unable to bind cocaine (DAT–Ci mice), showing complete lack of interest in cocaine self-administration (6,49,50). It was, nonetheless. somewhat surprising to find that drug-naïve DAT-AAA mice performed similar to WT mice in a naïve acquisition experiment using 1.0 mg/kg/infusion cocaine (Figure S1). This indicates that DAT-AAA mice can recognize the cocaine dose used as reinforcer in the active nose-poke hole versus no reinforcer in the inactive. Indeed, this is also supported by our in vivo microdialysis experiments demonstrating that cocaine did cause a modest increase in striatal DA levels in DAT-AAA mice, yet the response was significantly reduced compared to that seen in WT mice.

In conclusion, we present a mouse model with unique pre- and post-synaptic adaptations of the DA system that occur as a consequence of disrupting the ability of DAT to bind PDZ domain scaffold proteins to its C-terminus and that show interesting resemblance to the changes seen in the addicted brain. The mouse model might accordingly represent an important basis for further investigating the still poorly understood link between molecular alterations, changed DA homeostasis and compulsive drug or food intake.

## Materials and Methods

### Mice

DAT-AAA mice were generated as previously described (27). Adult male mice were acclimatized to the animal facilities, where experiments were conducted, for at least 1 week prior to experiments in group-housed (Macrolon type III cages) and kept on a 12-h light/dark cycle in a temperature and humidity-controlled room. Food and water were available ad libitum. All experiments were conducted during the light phase (8:00 a. m.–6:00 p.m.) conducted in accordance with guidelines from the Animal Experimentation Inspectorate, Denmark or in compliance with the Italian Ministry of Health (DL 116/92; DL 111/94-B) and European Community (86/609/EEC) directives regulating animal research. All efforts were made to minimize animal suffering and to reduce the number of animals used.

### Drugs

Cocaine obtained from HS Pharmacy (Copenhagen, Denmark) and quinpirole obtained from (Sigma Aldrich, Milan, Italy) were dissolved in 0.9% saline and prepared immediately before use.

### Super-resolution microscopy

Perfused brains were sliced at 10 µm on a cryostat (Leica CM2050 S) and immediately fused to 18 mm glass coverslips (#1.5) that had been treated with 3-aminopropyl-triethoxysilane (Sigma 440140). Samples were washed with PBS 3x 5 min, followed by a combination of glycine (20 mM) and NH_4_Cl (50 mM) in PBS 2x 15 min Next, they were washed with PBS 2x 5 min, and subjected to trisodium citrate (Sigma) (10 mM) in distilled water (pH 6.0, heated to 80°) for 30 min. Samples were again washed with PBS 4 x 5 min and then subject to blocking and permeabilization with 5% donkey serum, 1 % BSA, 0.3% Triton X-100 (Sigma) in PBS. Primary antibodies, in blocking and permeabilization buffer, were applied overnight at 4°. DAT MAB 369 was used to label DAT (1:200 dilution). VMAT2 was labeled with antibody graciously provided by Dr. Gary W. Miller (Columbia University, New York) (1:4000 dilution)) (51). The next day samples were washed with blocking and permeabilization buffer for 5 min, then 90-min increments. Secondary antibodies in blocking and permeabilization buffer were applied overnight at 4°. These were applied at concentrations of 10 µg/mL, were bought unlabeled from Jackson ImmunoResearch and conjugated to Alexa 647 for DAT or to CF568 for VMAT2. The next day the samples were washed with blocking and permeabilization buffer for 5 min, 5 mins, then 90-minute increments, followed by PBS for 2x 10 min, PFA (Electron Microscopy Supplies) (3%) for 15 min, a combination of glycine (20 mM) and NH_4_Cl (50 mM) in PBS 2x 15 min, and finally stored in PBS at 4° until imaging.

The dSTORM imaging was performed on an ECLIPSE Ti-E epifluorescence/Total Internal Reflection Fluorescence (TIRF) microscope (NIKON). Utilizing lasers at 405 nm, 561 nm and 647 nm, the sample was imaged through a 1.49 NA x100 apochromat TIRF oil immersion objective and a dichroic mirror with the range 350–412, 485– 490, 558–564, and 637–660 nm (97,335 QUAD C-NSTORM C156921). The emitted light was further filtered by a 561 nm longpass filter (Edge Basic, F76-561, AHF). Data was gathered by an EM-CCD camera (Andor iXon3 897). The imaging was performed with an imaging buffer containing 10% (w/v) glucose, 1% (v/v) ß-mercaptoethanol, 50 mM Tris-HCl (pH 8), 10 mM NaCl, 34 μg ml^−1^ catalase, 28 μg ml^−1^ glucose oxidase and 2 mM cyclooctatetraene. Data were collected for 20,000 frames per emission laser, alternating every frame between 561nm and 647 nm laser. Each frame was 16 ms. Localization data were obtained and refined through an automatic pipeline. First, localization data were obtained through ThunderSTORM (31). Raw images were subjected to a wavelet filter (B-Spline) with B-Spline order of 3 and B-Spline scale of 2. The method of molecular detection was non-maximum suppression with a peak intensity threshold of 0.9*std(Wave.F1) and a dilation radius of 3 pixels. The resulting data were drift corrected through a Matlab script for redundant cross correlation (52). The localizations were filtered for having an uncertainty value less than 25 nm and then localizations found within 15 nm and 3 frames were merged together.

Presynaptic sites were found according to the method defined in (29). In brief, the localization data were combined for the VMAT2 signal and the DAT signal for each image. The data were subject to a Voronoi tessellation and shapes larger than a certain size were removed (30). The remaining shapes were merged together, so that those that shared a border were now one shape. Following this merging, small areas were removed. The remaining shapes were smoothed around their edges, and used as masks for the dSTORM data, with each shape outlining a likely presynaptic varicosity.

Two different cluster metrics were used for this analysis, each utilizing DBSCAN (32). To compare cluster size, DBSCAN was applied to each varicosity with a search radius of 15 nm and a number of points qualifiers of 3 localizations. From there the convex hull was obtained for the localizations in each cluster and the diameter found from approximating the cluster as a circle. The fraction of the clusters with a radius of 75 nm or above was then found for each varicosity. To compare the presence of localizations in dense clusters, DBSCAN was applied to each varicosity with a search radius of 50 nm and a number of points qualifiers of 80 localizations. The fraction of localizations that satisfied these parameters per varicosity was compared.

### Stereotaxic surgery for microdialysis

Mice were premedicated with analgesic (Metacam 5 mg/kg s.c., Boehringer Ingelheim), anesthetized throughout the surgery with sevoflurane (4 vol.% sevoflurane in a mixture of 5% CO_2_ and 95% O_2_; Baxter) and mounted in a stereotaxic instrument (David Kopf Instruments, CA, USA). During surgery, the eyes of the animal were covered with a neutral eye ointment (Ophta, Rigshospitalets Apotek) for cornea protection. Prior to surgery, DAT-AAA and WT mice were treated with analgesic (Metacam 5 mg/kg s.c., Boehringer Ingelheim). The middle scalp was removed, and the skull was gently cleaned with aCSF containing 147mM NaCl, 4mM KCl and 2.3mM CaCl_2_, adjusted to pH 6.5. A small hole (d=1.2mm) was drilled, thus allowing intracerebral insertion of a microdialysis guide cannula (CMA7 guide cannula, CMA/Microdialysis AB) into the right dorsal striatum using the following coordinates according to the stereotaxic atlas of Franklin and Paxinos, 1997 (53): anteroposterior +0.5mm, mediolateral −2.0mm relative to bregma, and dorsoventral −3.0mm relative to skull surface. The microdialysis probe was secured to the skull with an anchor screw and dental acrylic cement (Dentalon Plus, AngThós AB, Lidingo, Sweden GC). Animals were then single-housed and allowed to recover for 24 hours following surgery.

### Total tissue contents of DA and metabolites

For determination of total tissue contents of DA and metabolites, mice were sacrificed by cervical dislocation and their brains were immediately taken out and briefly placed on parafilm covered pulverized dry ice, before striatal tissue was dissected out under microscope (54,55). Dissected striatal tissue of adult DAT-AAA mice and WT littermates were asweighed and homogenized in 500μl 0.1 N perchloric acid saturated with 5% Na_2_EDTA, and samples were quickly placed on crushed ice. Homogenates were then centrifuged at 14,000*g* for 20 min at 4°C. Supernatants were filtered through a 0.22μm cellulose acetate microfilter (13CP020AN Advantec, Frisenette ApS, Denmark), and 10μL was subsequently loaded on the HPLC-system for further analysis.

### Microdialysis procedure

Microdialysis disposables were obtained from CMA/Microdialysis AB (Solna, Sweden), or AgnThós AB (Lidingo, Sweden). Following recovery from surgery, a CMA/7 microdialysis probe (6,000 Dalton cut-off, CMA/Microdialysis AB; 2 mm dialysis membrane) was under sevoflurane (4 vol.% sevoflurane in a mixture of 5% CO_2_ and 95% O_2_; Baxter) anesthesia inserted into the guide cannula (56). The microdialysis probe was connected to a microinjection pump by a dual-channel swivel, allowing the conscious animals to move freely during the entire experiment in a bowl (Instech Laboratories, PA, USA). The 2 mm microdialysis probe was perfused at a flowrate of 2.0 μl/min for one hour with aCSF while the animals habituated to baseline. After habituation, the flowrate was switched to 1.8 μl/min and a 20-minute sampling regimen was initiated and maintained throughout the experiment using an automated refrigerated fraction collector (CMA 470). Three samples to establish DA baseline levels before the animals were injected with cocaine (30mg/kg s.c.) or saline, and dialysate fractions were collected for 140 min post-treatment. All dialysate fractions were analyzed immediately after collection using reversed-phase HPLC coupled to electrochemical detection. After termination of the microdialysis experiments, mice were sacrificed by cervical dislocation and brains were quickly removed for probe verification. The brains were cut on a cryostat (Shandon Cryotom, Pittsburgh, Pennsylvania, USA), and correct placement of probes was histologically verified under microscope.

### HPLC analysis of monoamines and their metabolites

Briefly, concentrations of DA, 3,4-dihydroxyphenylacetic acid (DOPAC), homovanillic acid (HVA), 5-hydroxyindoleacetic acid (5-HIAA), and serotonin (5-HT) in the tissue homogenates and of DA, DOPAC and HVA in the microdialysis samples were assessed by reversed-phase HPLC coupled to electrochemical detection. The HPLC system consisted of a HPLC pump (LC-20AD, Shimadzu, Kyoto, Japan), a degasser (LC-27A, Waters, Denmark), a Waters refrigerated microsampler (SIL-20ACHT, Shimadzu), an amperometric detector (Antec Decade II, Antec, Leiden, The Netherlands) and a computerized data acquisition system (LC Solution version 1.25, Shimadzu, Kyoto, Japan). The electrochemical detector cell was equipped with a glassy carbon electrode operating at +0.7V vs. Ag/AgCl reference electrode. Samples were injected onto a Prodigy C18 column (100Å∼2 mm I.D., 3-µm particle size, Phenomenex, Torrance, CA, USA). In the tissue homogenates, the monoamines and their metabolites were separated with a mobile phase consisting of 93% of 94.2 mM NaH_2_PO_4_, 0.98 mM octanesulfonic acid, 0.06 mM Na_2_ EDTA, adjusted to pH 3.7 with 1 M phosphoric acid and 7% acetonitrile (v/v) at a flow rate of 0.25ml/min, and detected by a glassy carbon electrode operating at +700mV vs. Ag/AgCl reference electrode. The output was recorded and the area under the curve for each peak calculated with LC Solution software by Shimadzu (Kyoto, Japan). Tissue concentrations of DA, DOPAC, HVA, 5-HIAA and 5-HT in the samples were estimated using a reference solution containing 0.5 or 1 pmol/sample of all compounds investigated. Concentration of monoamines and metabolites in the homogenate samples were extracted by their area under the curve (AUC) relative to AUC for the reference compounds, and tissue concentrations were determined relative to the weight of the tissue (μg/g tissue). Reference solutions were quantified before each set of homogenates. The limit of detection (at signal-to-noise-ratio 3) for DA was 7 fmol/20 μl. In the microdialysis samples, DA and its metabolites were separated with a mobile phase consisting of 55 mM sodium acetate, 1 mM octanesulfonic acid, 0.1 mM Na_2_EDTA and 8% acetonitrile, adjusted to pH 3.2 with 0.1 M acetic acid. 10 μl of the samples was injected and the flow rate was 0.25 mL/min.

### Fast scan cyclic voltammetry

Fast scan cyclic voltammetry experiments were performed as described (9). Briefly, mice were anaesthetized with isoflourane and decapitated. The brain was sectioned in cold carboxygenated artificial cerebrospinal fluid (aCSF) (125 mM NaCl, 2.5 mM KCl, 0.3 mM KH_2_PO_4_, 26 mM NaHCO_3_, 2.4 mM CaCl_2_, 10 mM D-glucose, 1.3 mM MgSO_4_) on a VT1000S vibrating microtome (Leica Microsystems, Nussloch, Germany) at a thickness of 300 µm. Coronal slices containing the dorsal striatum were allowed to recover for at least one hour at room temperature in carboxygenated aCSF. For recordings, slices were superfused with 32°C carboxygenated aCSF at a flow rate of 1ml/min. Experimental recordings started 20 min after transfer to the slice chamber. Carbon fiber electrodes (5 µm, Goodfellow, Huntingdon, England) were made as previously described (57,58). The carbon fibers were trimmed with a scalpel to 80-120 μm under a microscope (Nikon) A carbon 301 fiber microelectrode was inserted into the slice and a twisted bipolar stimulating electrode (Plastics One, Roanoke, VA) was placed on the surface of the brain slice ∼200 μm away. The potential of the working electrode was held at −0.4 V and scanned to +1.3 V and back at 300 V/s. Axonal DA release in the striatum was evoked by a single biphasic electrical pulse (1 ms long, 400 μA) every 2 min through a stimulus isolator (AM-system, Carlsborg, WA).

Data were filtered to reduce noise. Oxidation and reduction peaks were observed at ∼ +0.65 V and −0.2 V (vs. Ag/AgCl reference) identifying DA as the released chemical. Electrodes were calibrated in a flow injection system using 1 μM DA (Sigma Aldrich, Milan, Italy)

Data were normalized to the first 5 recordings (10 min) of their respective control period and graphically plotted against time (means ± SEM). Data analysis was performed using Demon Voltammetry software described (Yorgason et al.,2011). Briefly, computations were based on user-defined positions on current traces for baseline (Pre-Stim cursor), peak (Peak Cursor) and return to baseline (Post-Stim cursor) positions. Half-life values were determined from exponential fit curves based on Peak cursor and Post-Stim cursor positions using a least square constrained exponential fit algorithm (National Instruments, Milan, Italy).

### Autoradiography

For receptor autoradiography, DAT-AAA and WT mice were decapitated by cervical dislocation; brains were immediately taken out and frozen on dry ice. Brains were then stored at –80°C until sectioning. Using a cryostat (Shandon Cyrotome SME), 15μm coronal sections were cut to acquire striatal (1.54 mm to 0.74 mm relative to bregma), as well as VTA/SN (−3.08 mm to −3.64 mm relative to bregma) slices for D_1_R and D_2_R autoradiography. The sections were thaw-mounted onto glass slides (Superfrost Plus microscope, Menzel-Gläser, Germany), and allowed to dry before storage at −80 °C. While sectioning, orientation in the coronal plane was maintained by referring to a stereotaxic atlas of the mouse brain (53).

For detection of the D_1_R with [^3^H]SCH23390, glass slides with mounted tissue sections were incubated in pre-incubation buffer (50mM TRIS base, 120mM NaCl, 5mM KCl, 2mM CaCl_2_, 1mM MgCl_2_ and 20nM MDL-100907) at 0°C for 15 min. Application of the 5-HT_2A_ antagonist, MDL-100907 (H. Lundbeck, DK), was done to overcome the D_1_R radioligand ([^3^H]SCH23390) affinity for 5-HT_2A_ receptors. Sections were then incubated with a D_1_R antagonist for 1h at 4°C in a solution equivalent to the pre-incubation buffer with the addition of 1nM [^3^H]SCH23390 (Perkin Elmer, USA, 85 Ci/mmol). Non-specific binding was determined by incubating sections under the same conditions with excess amount of the D_1_R, D_2_R, and 5-HT_2_ receptor antagonist cis-(*Z*)-flupentixol (10μM, H. Lundbeck, Denmark). Sections were then washed for 2 x 5 min in pre-incubation buffer at 0°C and rinsed in demineralized water. Sections were allowed to dry in a flow bench overnight before they were exposed to Fuji BAS-TR2040 Phosphor imaging plates. Films were then exposed with standard [^3^H]-polymer-microscales (Batch 19, Amersham Biosciences, Denmark) for 17 days in autoradiography cassettes at room temperature prior to analysis.

For detection of the D_2_R with [^3^H]-raclopride, *g*lass slides with mounted tissue sections were incubated in pre-incubation buffer (50mM TRIS base, 120 mM NaCl, 5 mM KCl, 2 mM CaCl_2_, 1mM MgCl_2_) at 0 °C for 15 min. Sections were then incubated with a D_2_R antagonist for 1h at 4°C in a solution equivalent to the pre-incubation buffer with the addition of 4nM [^3^H]raclopride (Perkin Elmer, USA, 60.1 Ci/mmol). Non-specific binding was determined by incubating sections under the same conditions with excess amount of the D_2_R, 5-HT_2_R, and α_1_R antagonist sertindole (10μM, H. Lundbeck, DK). Incubation was followed by a washing phase for 2 x 30 min in pre-incubation buffer at 0°C in buffer. Sections were allowed to dry in the flow bench overnight before they were exposed to Fuji BAS-TR2040 Phosphor imaging plates. Films were then exposed with standard [^3^H]-polymer-microscales (Batch 19, Amersham Biosciences, Denmark) for 17 days in autoradiography cassettes at room temperature prior to analysis.

The Fuji BAS-TR2040 Phosphor imaging plates were read in a STARION FLA-9000 image scanner (Fujifilm Life Science, USA). The emitted radiation stored in the imaging plates was converted to a digital image. Densitometric image analysis of autoradiograms was achieved using WCIF ImageJ software (Ver. 1.37a, NIH, USA) to quantify D_1_R and D_2_R levels. Regions of interest (dorsal striatum, ventral striatum and ventral midbrain) were selected for each section, and the mean pixel density in the selected regions was measured. Image density in the selected regions was expressed in arbitrary units as a grey-scale level with a value designated each pixel according to its density. Quantification of receptor binding was achieved by calibrating the pixel density to decay-corrected [^3^H]-polymer-microscale activity levels. The calibration allowed for quantification of receptor levels in Bq per mg tissue instead of arbitrary pixel densities. Levels of receptor binding were measured bilaterally in the dorsal and ventral striatum. Values from both sides (right and left) were averaged for each section and animal. For better visualization, and to acquire higher anatomical resolution of the autoradiograms, exposure to tritium-sensitive films (KODAK Biomax MR, Amersham Biosciences, Denmark) at −20°C for 85 days was achieved succeeding phospho-image exposure. Films were developed in a Kodak GBX film developer and fixer (Sigma Aldrich, Denmark).

### Radioligand binding assay

Adult mice were sacrificed by decapitation and brains were rapidly removed. Striata from DAT-AAA and wildtype mice were dissected from coronal slices and homogenized in ice-cold homogenization buffer (50 mM Tris-HCl, 1 mM EDTA, pH 7.4) using a motor-driven teflon pestle with 10 even strokes at 800 rpm. Homogenate was centrifuged at 16,000 x g at 4°C for 30 min to isolate the membrane fraction. Membranes were re-suspended in ice-cold binding buffer (25 mM HEPES, 120 mM NaCl, 5 mM KCl, 1.2 mM CaCl_2_, 1.2 mM MgSO_4_, 1 mM L-ascorbic acid, 5 mM D-glucose, pH 7.4) followed by assessment of protein concentration using BCA™ Protein Assay kit (Pierce). Saturation binding experiments were performed on striatal membrane preparations using various concentrations of ^3^H-raclopride, a D_2_R antagonist (71.3 Ci/mmol, Perkin Elmer, USA). For determination of non-specific binding, quinpirole hydrochloride (0.3 mM), a D_2_R agonist was included. Membrane suspensions (final incubation, 0.3-0.4 mg protein/ml) were mixed with ^3^H-raclopride in binding buffer (a total volume of 500µl) containing 25 mM HEPES pH 7.4, 120 mM NaCl, 5 mM KCl, 1.2 mM CaCl_2_, 1.2 mM MgSO_4_ for 1h at room temperature with constant shaking. Membranes were then immediately loaded on glass microfiber filters (GF/C Whatman®), rinsed 2 x 8 ml in ice-cold binding buffer and allowed to air dry. Scintillation fluid was added and filters were agitated for 1h and counted in Wallac Tri-Lux β-scintillation counter (Perkin Elmer). Binding data were analysed by non-linear regression analysis assuming one-site binding (GraphPad Prism 5.0).

### Immunohistochemistry

Adult mice were anaesthetized and transcardially perfused with heparinized phosphate buffered saline (PBS) followed by 4% paraformaldehyde in 0.1 M PBS. The brains were isolated, post-fixated in paraformaldehyde for 24h and transferred to 20% sucrose for cryoprotection. Brains were then rapidly frozen on powdered dry ice and kept at −80°C until further processing. Subsequently, coronal sections (40 μm) from striatum and midbrain were generated and stored in anti-freeze solution at −20°C. For bright-field immunohistochemistry, a standard peroxidase-based method using 3, 3’-diaminobenzidine (DAB) was applied. Free-floating tissue sections were rinsed in PBS three times followed by preincubation with 5% rabbit serum in PBS containing 0.3% Triton X-100 and 1% bovine serum albumin (BSA). Sections were then incubated with rat polyclonal D_1_R antibody (1:1000, Sigma-Aldrich, USA) at 4°C overnight. On the second day, sections were rinsed and incubated with biotinylated rabbit anti-rat IgG for 1h (1:200, DAKO Cytomation A/S, Denmark). Following avidin-biotin-peroxidase complex (Vector laboratories, USA) incubation, peroxidase was visualized using DAB (0.5 mg/ml) and 0.01% H_2_O_2_ treatment. After additional washing, sections were mounted on SuperFrost slides (Menzel-Gläser, Germany), air-dried overnight and finally cover-slipped using Pertex (Histolab, Sweden).

### Operant Behavior

Intravenous self-administration equipment, training, and evaluation procedures were previously described (59). In short, the operant chambers (Med Associates, USA) contained two nose-poke holes 10 mm above the grid floor, both equipped with photocells and a discriminative cue light, positioned on either side of a small dish shaped plate into which liquid food could be delivered. Responding in the right hole resulted in delivery of a reinforcer and illumination of the cue light for 20 s, during which additional responses were counted but had no scheduled consequences (timeout responses).

### Catheter implantation surgery and maintenance

Under isoflurane vapor anesthesia, a catheter (SILASTIC tubing; 0.18-mm inner diameter, 0.41-mm outer diameter, CamCaths, UK) was inserted 1.2 cm into the right or left jugular vein and anchored to the vein with sutures. The catheter ran subcutaneously to the base located above the midscapular region. During the subsequent 7 days of postsurgical recovery, 0.02 ml of 0.9% saline containing heparin (30 U/ml; SAD, Denmark) and antibiotic (cefazolin, 50 mg/ml; Hexal, Germany) was infused daily through the catheter to prevent clotting and infection. Catheter patency was confirmed 7 days after surgery and after completion of each experimental phase by the loss of muscle tone and clear signs of anesthesia within 3 s after infusion of 0.02– 0.03 ml ketamine (15 mg/ml; Pfizer, Denmark) and midazolam (0.75 mg/ml; Matrix Pharmaceuticals, Denmark) in saline.

### Liquid food self-administration under a FR schedule

A separate set of experimentally naïve mice was used for self-administration of a nondrug reinforcer under a FR schedule. The mice were mildly food deprived before the first presentation of liquid food (5 ml of Nutridrink high energy drink, vanilla flavor, Nutricia A/S, Allerød, Denmark) in the operant chamber (i.e., ad libitum dry food in the home cage was removed 18–20 h before the session, water remained available). When ≥1.5 ml of the 5 ml available was consumed per 2-h session, mice were placed in the operant chamber with one active and one inactive nose-poke hole for daily 2-h sessions similar to cocaine FR 1 self-administration. Acquisition lasted for at least five consecutive sessions and until criteria were met (≥20 reinforcers earned, with ≤20% variation over two consecutive sessions and ≥70% responses in the active hole). Subsequently, water was substituted for at least three sessions and until responding was extinguished to <80% of food-maintained responding. Then, a range of liquid food dilutions (Nutri drink: water; 3%, 10%, 32%, and 100%) was presented according to a Latin-square design, determined twice in each mouse.

### Cocaine self-administration and liquid food-maintained behavior under a PR schedule

After the FR 1 schedule, mice proceeded with cocaine self-administration under a PR schedule of reinforcement. Mice that had self-administered liquid food under a FR 1 schedule proceeded in a similar PR schedule with liquid food as the reinforcer. After stable responding under the FR 1 schedule maintained by cocaine (1.0 mg/kg/infusion) or undiluted food, an FR 3 schedule was used as a transition from the FR 1 schedule before introducing the PR schedule. For the PR schedule, the starting ratio was 3, and then increased by 0.115 log units after each reinforcer delivery (i.e., 3, 4, 6, 7, 10, 13, 16, 21, 28, 36…). The breaking point was defined as the step value associated with the last completed ratio (i.e., number of reinforcers earned) after a 60-min limited hold (i.e., period with no reinforcer earned). If a breaking point was not reached within 6 h, the session was terminated to prevent health hazard, and the last reached ratio was used. After stable baseline was achieved (two consecutive sessions with breaking points >5 and with <20% variation), saline or water was substituted until responding extinguished to ≤50% of the baseline breaking point. Cocaine dose–effect curves (0.1, 0.32, 1.0, 3.2 mg/kg/infusion cocaine) and liquid food concentration–effect curves (0%, 3%, 10%, 32%, and 100% food in water) were determined according to a Latin-square design, with each dose tested for two or three consecutive sessions (i.e., if the breaking points reached in the two first determinations varied by >20%, a third determination was made).

### Drug-induced locomotion

Locomotor activity was measured in monitoring frames equipped with eight horizontal infrared light beams along the long axis of the frame placed 4.3 cm apart and 3.3 cm above the surface as previously described (27,60). Standard cages (macrolon type III; 37×21×17 cm) with a lining of fresh wood-chip bedding were placed in the monitoring frames and covered with Plexiglas tops with ventilation holes. The set-up was situated in a ventilated room. A computer program (MOTM, Ellegaard Systems, DK) recorded interruptions of the infrared light beams as counts of beam breaks in intervals of five min. Cages were cleaned between tests. Mice were injected with cocaine or saline i.p. 3-7 min before testing. The animals were then placed in a cage and the activity was measured for 1 hour.

### Statistics

All statistical analyses were performed using GraphPad Prism 8 (www.graphpad.com). Two-way ANOVA was used together with Holm-Sidak post-hoc test for analysis of DA and DA metabolite tissue content, food and cocaine self-administration and cocaine-induced locomotor activity data. A three-way repeated measures (RM) ANOVA was used together with Holm-Sidak post-hoc test for analysis of microdialysis data. DA receptor binding data (autoradiography) were analyzed by use of multiple t testing with Holm-Sidak correction while [^3^H]-Raclopride binding (B_max_) was analyzed by a one-sample t-test. FSCV data and super-resolution microscopy data were analyzed by unpaired t-tests or Wilcoxon matched-pairs signed rank test. Significance level was set at *P*<0.05 in all analyses.

## Data availability Statement

All data relating to this manuscript is contained in the manuscript.

## Acknowledgements

We thank Pia Elsman, Birgit Hansen, Saiy Kirsari, and Pernille Clausen for excellent technical assistance.

## Funding and additional information

The work was supported by the University of Copenhagen BioScaRT Program of Excellence (G.S., D.P.D.W., G.W., A.F. and U.G.), the National Institute of Health Grants P01 DA 12408 (UG), Independent Research Fund Denmark – Medical Sciences (U.G. 4004-00097B), Lundbeck Foundation R199-2015-2110 and R77-2010-6815 (UG and MR), the Novo Nordisk Foundation (NNF16OC0023104) (U.G.) and St. Petersburg State University, St. Petersburg, Russia (project ID: 51143531) (RRG).

## Supplementary figure legends

**Figure S1.**
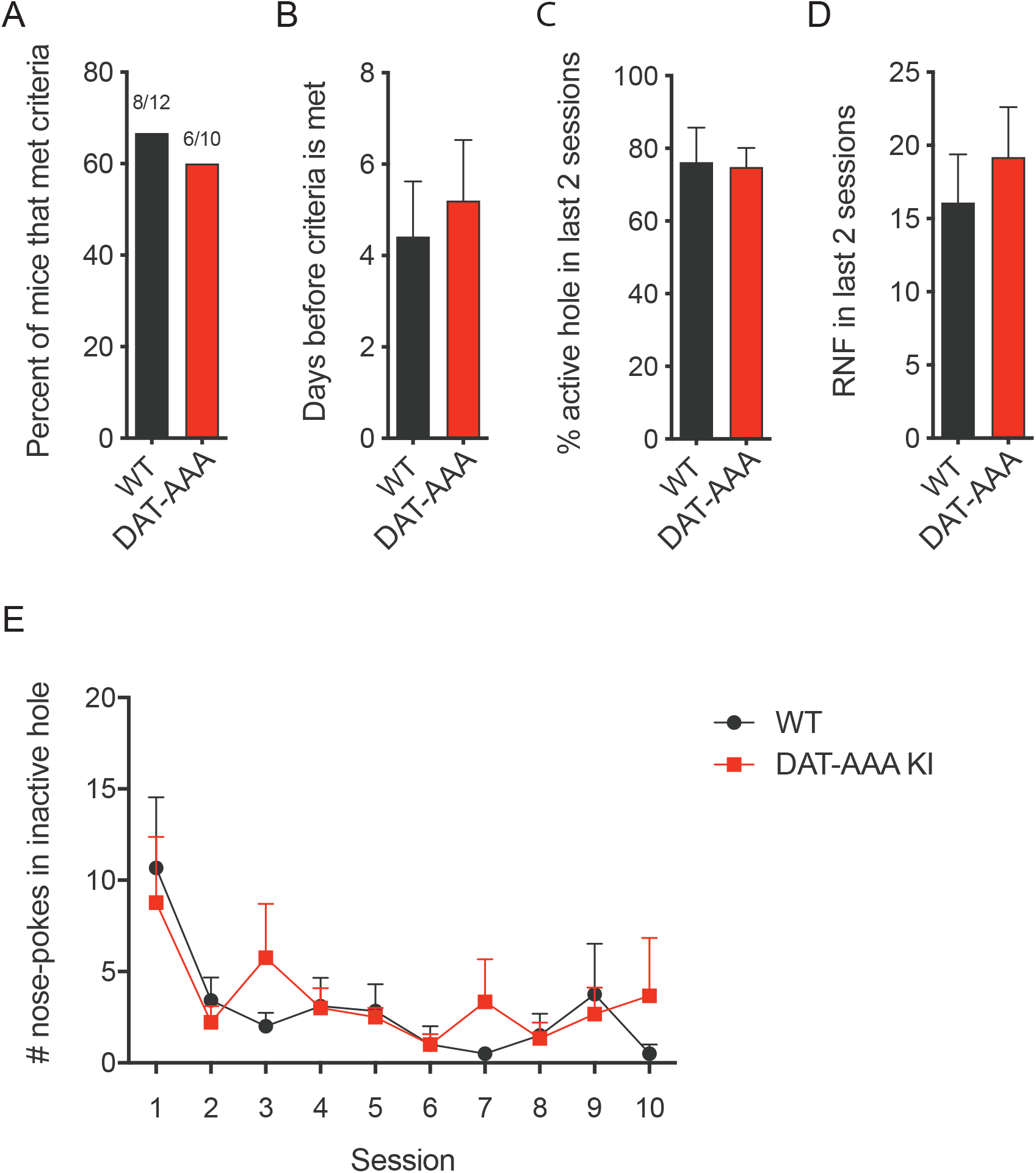
Operant naïve acquisition of 1.0 mg/kg/infusion cocaine. Operant naïve acquisition of 1.0 mg/kg/infusion showed no difference between DAT-AAA and WT mice in any of the criteria measured; *A*, Percent of mice that met criteria. *B*, Days before criteria was met. *C*, Percent nose pokes in the active nose poke hole at criteria. *D*, Number of reinforcers at criteria. *E*, Percent nose pokes in the active nose poke hole through-out all 10 sessions.

